# A three-dimensional *ex vivo* model recapitulates *in vivo* features and unravels increased drug resistance in childhood acute lymphoblastic leukemia

**DOI:** 10.1101/2024.12.13.628110

**Authors:** Magdalini Kanari, Iria Jimenez Garcia, Fabio D. Steffen, Lisa A. Krattiger, Charles Bataclan, Wangjie Liu, Benjamin R. Simona, Bart Deplancke, Olaia Naveiras, Martin Ehrbar, Beat Bornhauser, Jean-Pierre Bourquin

## Abstract

Acute lymphoblastic leukemia (ALL) preferentially localizes in the bone marrow (BM) and displays recurrent patterns of medullary and extra-medullary involvement. Leukemic cells exploit their niche for propagation and survive selective pressure by chemotherapy in the BM microenvironment, suggesting the existence of protective mechanisms. Here, we established a three-dimensional (3D) BM mimic with human mesenchymal stromal cells and endothelial cells that resemble vasculature-like structures to explore the interdependence of leukemic cells with their microenvironment. This model recapitulates recurrent topologic differences between B-cell and T-cell precursor ALL, whereby B-ALL interacts more closely with the mesenchymal compartment. Migration versatility was found to be associated with subtype, consistent with increased motility observed in T-ALL *in vivo*. Single-cell RNA signatures revealed similarities to profiles from *in vivo* patient derived xenografts, suggesting relevant states *ex vivo*. Furthermore, enhanced migration, adherence and cell cycle heterogeneity was visualized in our co-culture model. Finally, drug response profiling experiments in this 3D system reproduced established response patterns and indicated that drug resistant leukemic subpopulations may be detected more faithfully compared to information from two-dimensional models.

## Introduction

Acute lymphoblastic leukemia (ALL) is the most common pediatric cancer originating from the malignant transformation of lymphoid progenitor cells in the bone marrow (BM) with frequent involvement of medullary and extra-medullary sites.^1^ During leukemogenesis, the BM forms a protective ecosystem which contributes to the propagation and acquired chemoresistance of ALL cells leading to relapse.^2–4^ The non-hematopoietic compartment, including mesenchymal stromal cells (MSCs), osteo-lineage cells, endothelial cells (ECs), fibroblasts, chondrocytes and adipocytes, plays a substantial role in hematopoiesis and has been directly linked to the transformation of the BM into a tumor supporting niche.^5,6^ Increased vascularization, adhesive interactions with ECs, MSCs and osteoblasts through expression of CXCR4 (CXC motif chemokine receptor 4), VLA-4 (very late antigen-4) and CD44 (homing cell adhesion molecule) molecules as well as hypoxia induction are amongst common alterations in the leukemic BM.^7^ Fibroblast- driven extracellular matrix (ECM) remodeling, reduced bone density by impaired osteoblast function, fatty acid consumption and adipocytic employment for chemoresistance further sustain this leukemic niche.^8,9^ We have previously shown by using *in vivo* patient-derived xenograft (PDX) models that residual leukemia cells can survive chemotherapeutic stress in the BM microenvironment without clonal selection or additional mutations, suggesting the presence of protective cues in this niche.^10^ Nonetheless, these molecular signaling events have thus far remained elusive yet will represent candidates for interference to eradicate resistant populations.

Two-dimensional co-culturing systems support leukemic expansion but lack the hierarchical complexity and cell communication that occurs in the BM.^11^ In recent years, three-dimensional models have emerged as a promising tool for the study of leukemia to identify underlying interactions with its niche as well as to predict drug response more accurately by recapitulating the microenvironment protection.^12^ Different approaches have been implemented including scaffold-based systems like hydrogels, spheroids, organ-on-a-chip applications, or microfluidic devices.^13–22^ In all of the studies, the authors describe leukemia- stromal interactions, modulated gene expression and phenotypic plasticity as well as increased chemoresistance induced by supporting microenvironmental cues. Patient-derived MSC spheroids support the expansion of leukemic samples and validate the presence of a leukemic subpopulation that mimics minimal residual disease signatures.^14^ 3D microfluidic chips, resembling the *in vivo* morphology and cellular composition of the BM, have demonstrated dynamic interactions between cellular types and linked microenvironment-promoted quiescence to increased drug resistance.^13,15^ However, these systems are complex, difficult to reproduce at scale and lack general applicability for patient-derived leukemic models.

In this study, we developed a 3D BM model compatible with high-content screening in 96-well imaging plates. The three component system consists of human MSCs and ECs that form 3D blood vessel-like structures, readily providing a niche for culturing patient-derived ALL cells. We identified distinct ALL features in the presence of the supporting cells and identified topological patterns resembling *in vivo* localization. We performed single-cell RNA sequencing (sc-RNAseq) of all cellular compartments to identify molecular cues that underlay the leukemic-microenvironment interactions and phenotypically and functionally characterized 13 leukemic PDXs in this system. Finally, we investigated whether drug resistance is driven by the 3D support and compared it with two-dimensional (2D) response in order to identify potential enhanced predictive accuracy of drug testing.

## Results

### A 3D vasculature-like model for ALL patient-derived xenografts

To evaluate the role of the non-hematopoietic BM in the leukemic ecosystem, we developed a 3D model that mimics blood vessel-like structures, enabling functional studies of their interactions with leukemic cells. This was achieved by co-culturing stromal, endothelial and leukemic cells in 96-well imaging plates containing pre-cast, optically transparent synthetic hydrogels. These hydrogels are based PEG-peptide bioconjugates and contain adhesion and degradation motives supporting cell spreading, MMP-mediated gel degradation and formation of cell-produced ECM. Moreover, an in-depth surface density gradient allows the characterization of motility patterns for different cell types (Fig. 1A).^23–27^ The primary human BM MSCs are necessary for endothelial spreading, acting as a supporting scaffold (suppl. Fig.1A), while no “vascularization” occurs in their absence (suppl.Fig.1B). Interestingly, the hTERT-MSC (human telomerase reverse transcriptase-MSC) cell line is not capable of supporting this “vascularization”, highlighting the importance of using primary MSCs for a blood-vessel mimic (suppl.Fig.1C). 3D segmentation and characterization of the structures revealed consistent volumes and surfaces of network amongst the control, an MSC-HUVEC co-culture, and MSC-HUVEC-ALL co-cultures established with 10 PDXs, thus validating the robustness of the model across multiple conditions (Fig. 1B-D).

**Figure 1.**
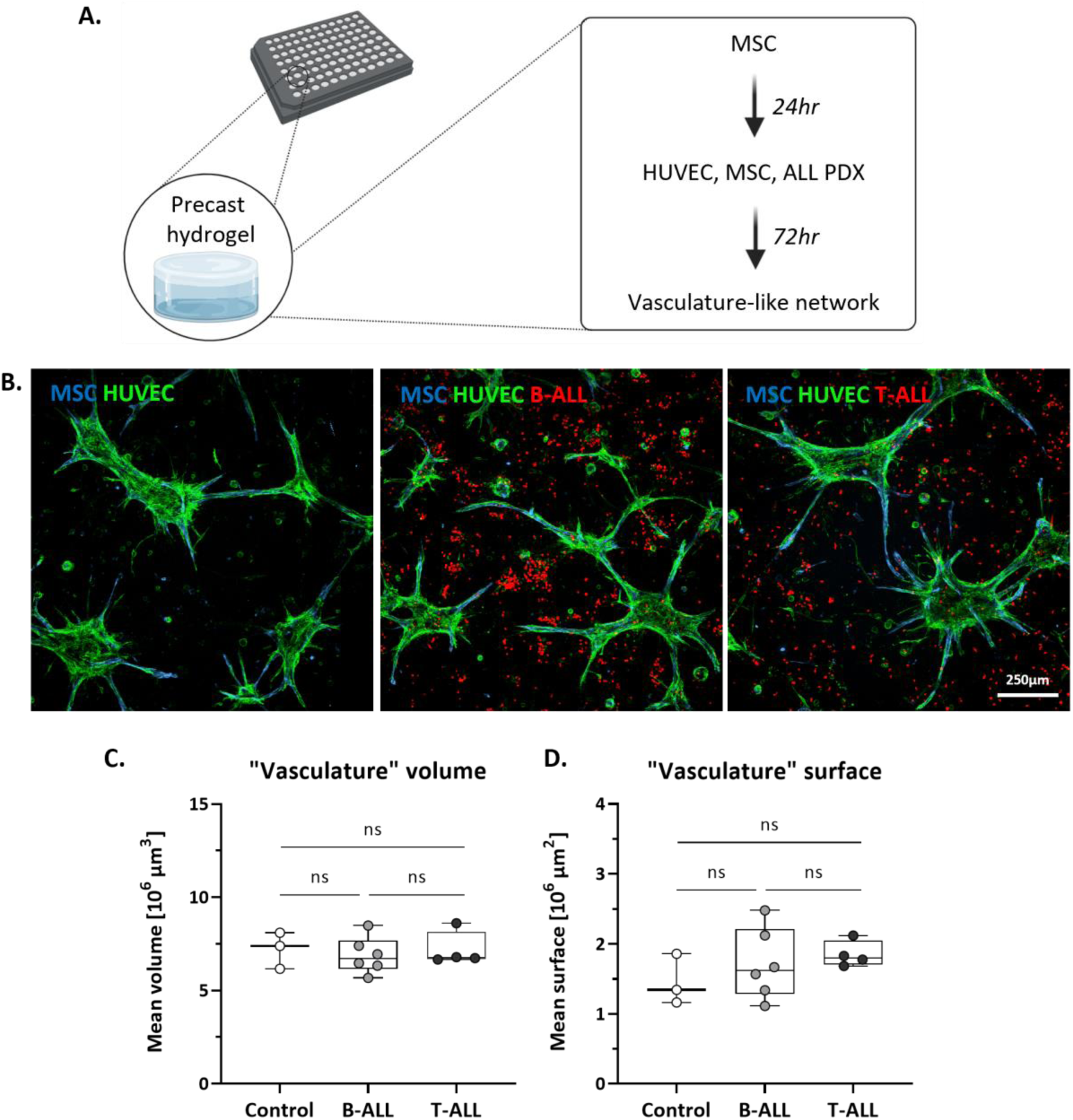
Description of the 3D vasculature-like model for leukemic PDXs. A. Experimental set-up. 3DProSeed® Hydrogel Well Plate was used for the establishment of the ex vivo model. Primary MSCs are cultured for 24 hours, followed by the addition of HUVECs, MSCs and ALL PDXs. B. Representative confocal fluorescence images of the co- cultures in maximum projection, z-stack with 10 μm distance (MSC: blue, HUVEC: green, ALL: red). From left to right: MSC-HUVEC control, MSC-HUVEC-B-ALL, MSC-HUVEC-T-ALL. C,D. The “vascularization” occurs in a similar manner in all 11 conditions tested (MSC-HUVEC controls, 10 PDXs co-cultures). Depicted are the mean volume and mean surface of the HUVECs (“vasculature”) in triplicates calculated by 3D segmentation. Controls were quantified in triplicates in three separate experiments (statistics: unpaired t test, significant p value <0.05).

### Leukemic cells exhibit enhanced motility upon immediate interaction with the supporting cells

To analyze the influence of the microenvironment on leukemic cell behavior, we performed timelapse confocal imaging directly after seeding for 30 hours with a timestep of 5 minutes, comparing 3D ALL mono- cultures and ALL co-cultures with MSCs and ECs. It is evident that the leukemic PDXs directly interact with the supporting cells as they form multi-cellular clusters over time. Single-cell distance analysis revealed enhanced leukemic aggregation over time, with decreasing ALL distances in a 12-hour course (Fig. 2A). Significantly higher speed and displacement in the XY plane was observed for the leukemic cells in co-culture compared to mono-culture, as measured in the first 6 hours of the timelapse (Fig. 2B,C). The movement of the ALL cells in the z-axis was also influenced by the presence of supporting cells. Figures 2D,E show the ALL cell vertical distribution over time. ALL cells in mono-culture remained concentrated around the median position, with decreasing standard deviation from 0 to 30 hours, indicating no substantial migration. In contrast, in co-culture we observed that the leukemic cells interacted with the supporting cells, changing their migratory phenotype and exhibiting a subpopulation with increased penetration capacity, as indicated by the increased values of delta median and standard deviation at time 30 compared to time 0.

**Figure 2.**
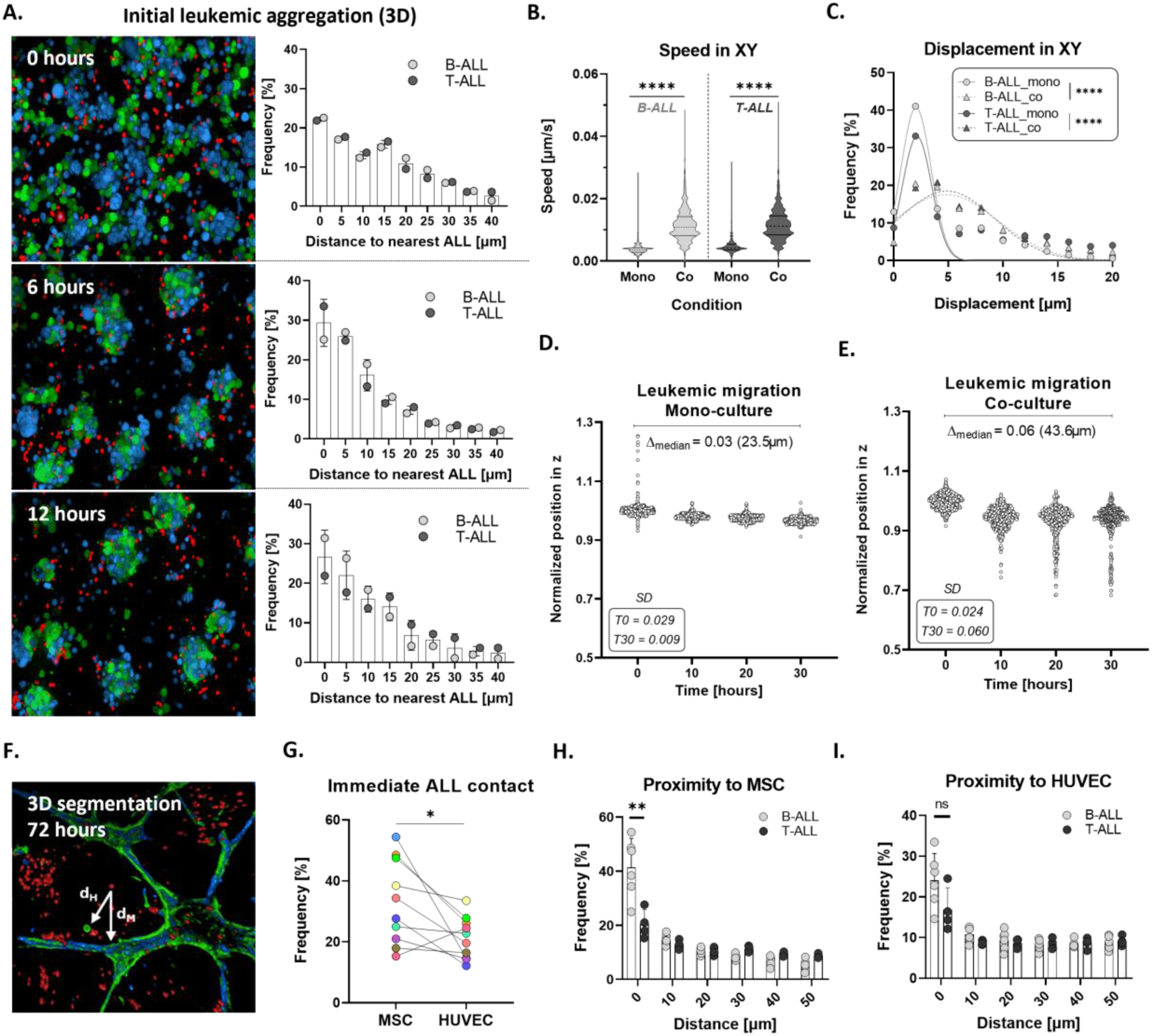
ALL PDX distinct phenotypes upon immediate interaction with microenvironment cells. A. Maximum projections of confocal images (z-step: 7.8μm) showing the immediate formation of multi-cellular clusters (MSC: blue, HUVEC: green, ALL: red). The leukemic cells follow the clustering of MSCs and HUVECs towards the network formation (12 hours, 5 minute interval). The corresponding bar plots show the decreased distances amongst the ALL cells (3D single-cell analysis, bin center = 5). B, C. XY speed and motility analysis of segmented ALL single cells (left: speed analysis, right: displacement analysis, bin center = 2, statistics: Kolmogorov-Smirnov unpaired t tests with p < 0.0001). D, E. Image-based single-cell segmentation and quantification of the leukemic distance to the bottom in z, normalization based on the median position of each timepoint. Data obtained by 30-hour live cell imaging with a 5 minute interval. Depicted is the Δ in z between the median position from timepoint 0 to 30 based on the normalized data (and actual values in μm), as well as the standard deviation (SD) at 0 (T0) and 30 (T30) hours. F. 3D segmentation of all cellular types at 72 hours enables proximity analysis, dH: minimum ALL distance to HUVEC, dM: minimum ALL distance to MSC. G. Percentage of ALL cells residing at a distance of <10μm from MSC and HUVEC (statistics: paired Wilcoxon signed rank test with p <0.05, triplicates for all conditions and 10 different PDXs as biological replicates). H, I. Comparison of topological ALL features based on the subtype (bin center = 10). B-ALL PDXs are in closer contact with both MSC and HUVEC compared to T-ALL PDXs. Statistically significant differences are shown only for the MSC positioning (statistics :unpaired t test with Welch’s correction, p value <0.05, triplicates for all conditions).

Furthermore, the positioning of the leukemic cells towards MSCs and HUVECs was evaluated after 72 hours, where the network formation is completed. 3D segmentation was performed and the minimum distance to the respective cell type (dH: ALL distance to HUVEC, dM: ALL distance to MSC) was calculated (Fig. 2F). Increased 3D proximity to MSCs compared to HUVECs was observed for the ten PDXs analyzed, while the localization towards the two cell types was not directly correlated (Fig. 2G). Proximity analysis accounting for the leukemic subtype revealed that while B-cell precursor ALL cells closely localized to stromal or endothelial cells, T-ALL cells were more versatile and did not preferentially localize to distinct cellular structures, which mimics the phenotypes that we described using a patient-derived *in vivo* xenograft model (Fig. 2H,I).^10^

### Multi-lineage differentiation of MSCs in the blood-vessel mimic maintains *in vivo* cell fate

To further elucidate the heterogeneity across the ALL PDXs and their related BM niches, we performed single-cell RNA sequencing (scRNA-seq) to decipher transcriptomic signatures driving these distinct phenotypes. The conditions included all the cell types cultured alone, an MSC-HUVEC control and 6 PDX leukemic co-cultures (MSC-HUVEC-ALL, referred as CO1-CO6) of different ALL subtypes. With the use of established cell surface markers (CD19, and CD7 for B- and T-ALL, THY-1 for MSC and CDH5 for HUVEC) for their identification, both supporting and leukemic cells were analyzed. Enhanced MSC differentiation was observed in the presence of HUVECs, both in the control and leukemic conditions, and well-established gene sets were used to identify these clusters (Fig. 3A).^3,28–34^ Notably, a polarization based on the known MSC-markers PDGFRA (platelet-derived growth factor receptor alpha) and CXCL12 (CXC motif chemokine 12) was observed, which is in line with reports on *in vivo* and human primary MSCs, with PDGFRA-MSCs exhibiting high multipotent capacity and CXCL12-MSCs preferentially committed to osteo- lineages (Fig. 3B).^35–38^ 9 subpopulations were identified including precursors of osteo-, adipo-, fibro- and chondro-lineages, with some characterized by the co-expression of genes related to more than one lineage, cycling MSCs with high MKI67 expression and more committed populations such as perivascular and smooth muscle cells (Fig. 3C, suppl.Fig.2A-C). Moreover, increased differentiation towards smooth muscle cells and pre-fibroblasts is observed, potentially due to their capability of producing vascular promoting and supporting factors such as the vascular endothelial growth factor (VEGF).^39,40^ A slight further enrichment in the fibro-lineage (5%) is detected for the leukemic conditions, possibly describing the transition into a cancer-associated fibroblastic phenotype that facilitates the homing of the ALL cells close to the network (Fig. 3D).^41^ Furthermore, gene set enrichment analysis (GSEA) highlighted the role of the MSCs on “vascularization”, as blood vessel morphogenesis and endothelium development are the most upregulated pathways in the HUVEC-containing conditions (Fig. 3E). Notably, when looking into the comparison of leukemic MSCs to MSCs only, pathways related to external encapsulating structure organization are enriched, while not in the MSC-HUVEC control, suggesting the re-structuring of the MSC surrounding microenvironment to facilitate leukemic development (suppl.Fig.2D,E). Finally, cell-cell communication network analysis revealed known supporting interactions between MSCs and ALL, with the highest probability being through CD74 and CD44 signaling (suppl.Fig.2F).

**Figure 3.**
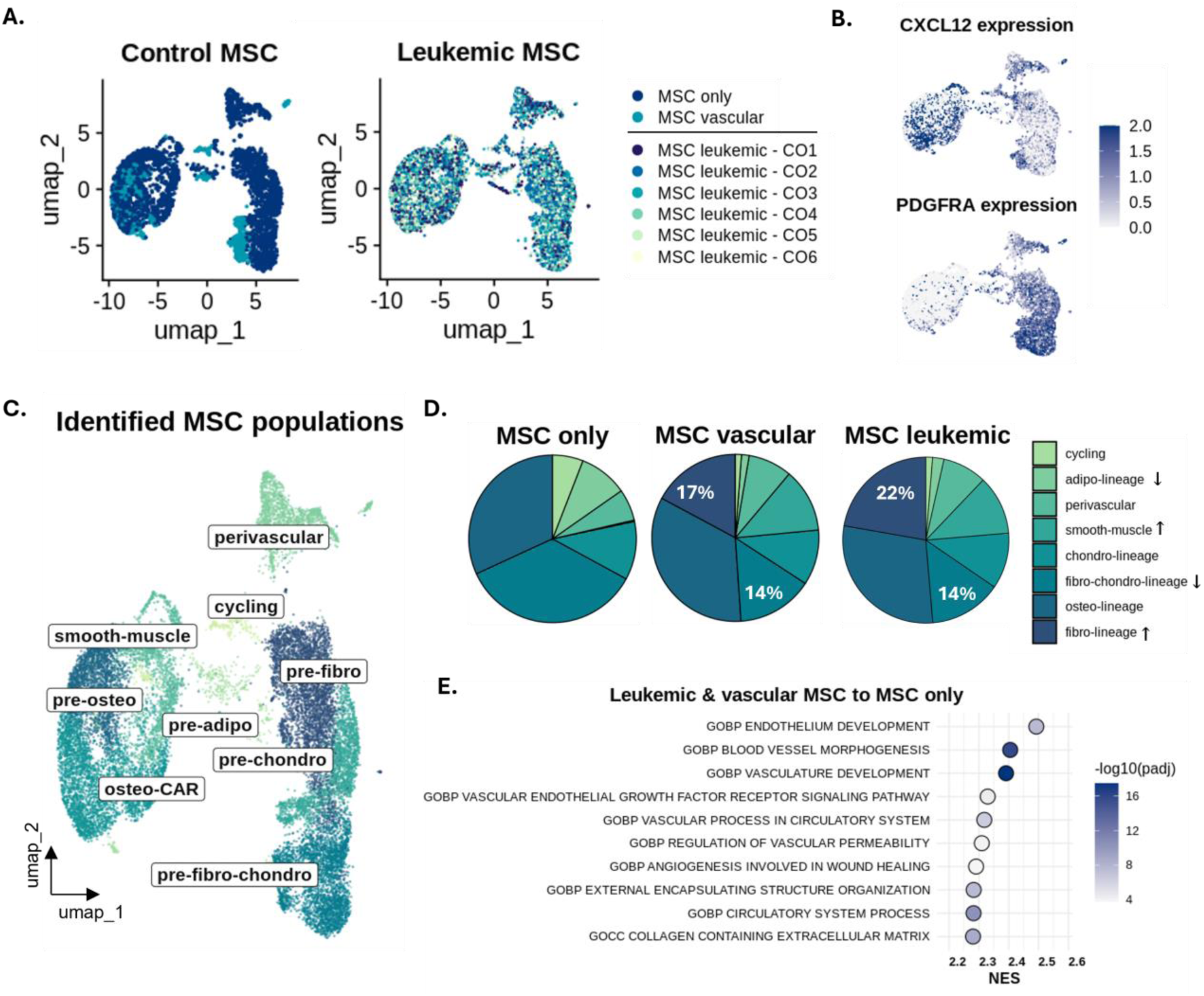
MSC multi-lineage differentiation in 3D resembles in vivo and human data. A. UMAP projections showing MSC population overlap based on the culturing condition, left plot depicting the two controls, i.e. MSCs cultured alone (MSC only) and MSCs cultured with HUVECs (MSC vascular), while right plot depicting the MSCs cultured with HUVECs and ALLs (MSC leukemic, CO1-CO6 indicating the six different ALL PDXs). B. Feature plots depicting the expression of CXCL12 (top) and PDGFRA (bottom) across the MSC populations. C. Identification of multi-lineage MSC subpopulations, projected in UMAP, upon coculturing in 3D. D. Proportions of the subpopulations based on the condition, with the increased population percentages in HUVEC-containing conditions in white. Arrows indicating increase or decrease upon co-culture E. GSEA dot plot showing top 10 enriched pathways in vascular and leukemic MSC compared to MSC only.

### GSEA reveals EC-induced ECM reorganization in the presence of leukemic cells

On the other hand, ECs exhibit less complexity but still consist of distinct subpopulations including a mesenchymal-like, a pre-vascular and a vascular cluster (Fig. 4A-C), with a notable transition to the vascular-like phenotype observed in all co-cultures (Fig. 4D). ^28,42,43^ Another significant observation is the enrichment of the extracellular matrix degradation pathway, and other ECM-related ones, exclusively in the leukemic conditions, while in the control only enhanced proliferation is observed (Fig. 4E, suppl.Fig.2G). This could indicate that endothelial cells degrade and re-organize the ECM upon leukemic infiltration in order to create a favorable niche, as they are already known regulators of ECM dynamics.^26^

**Figure 4.**
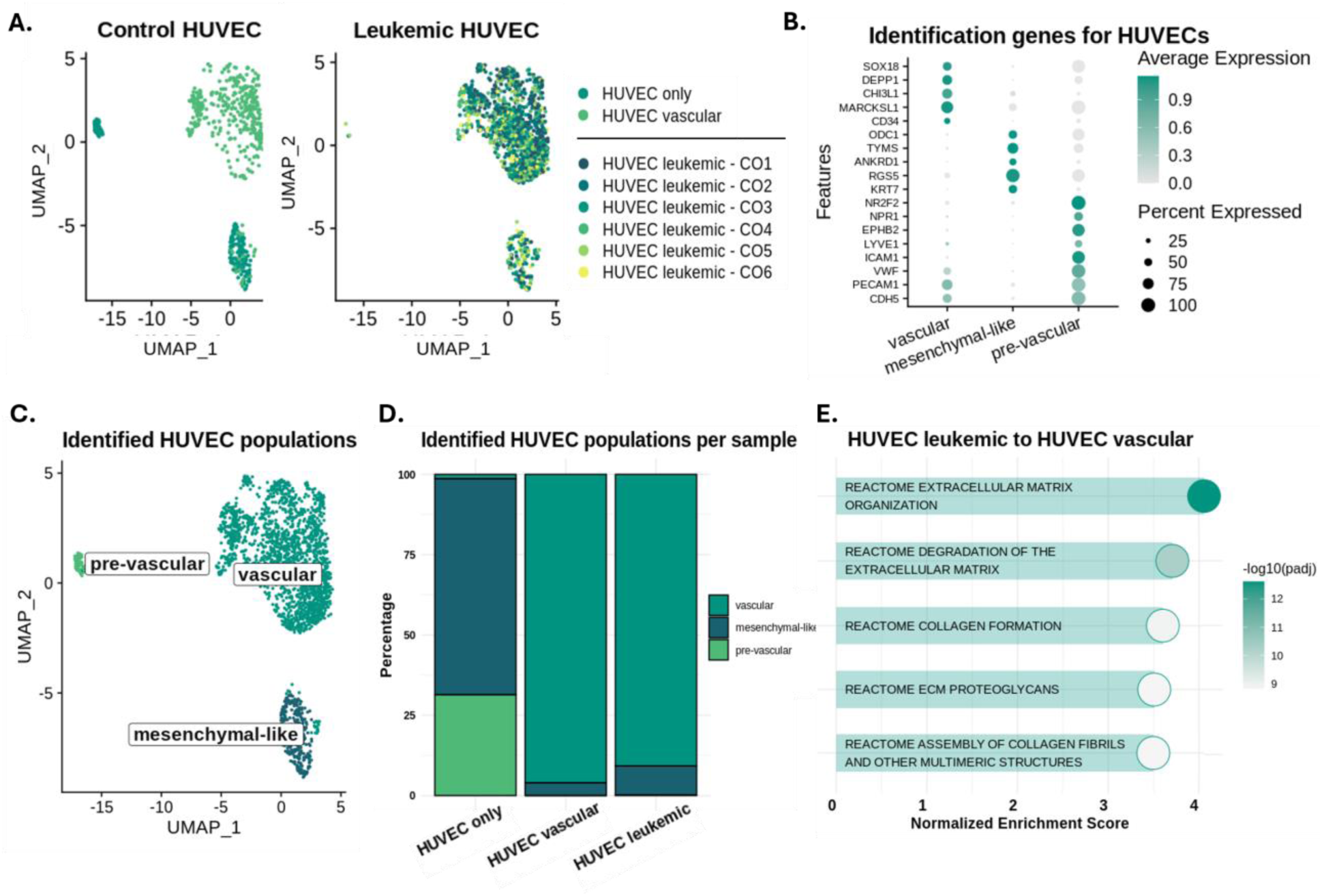
Endothelial cells re-organize the microenvironment in the leukemic conditions. A. UMAP projections showing HUVEC population overlap based on the culturing condition, left plot depicting the two controls, i.e. HUVECs cultured alone (HUVEC only) and HUVEC cultured with MSCs (HUVEC vascular), while right plot depicting the HUVECs cultured with MSCs and ALLs (HUVEC leukemic, CO1-CO6 indicating the six different ALL PDXs). B. Dot plot depicting the expression of genes characterizing each endothelial cell cluster. C. UMAP projection with the corresponding HUVEC subpopulations. D. Stacked bar plot depicting the percentage of the HUVEC subpopulations per condition. E. GSEA dot plot showing degradation of the ECM amongst the top 5 pathways enriched in leukemic HUVECs.

### Leukemic gene expression in the 3D microenvironment mirrors signatures detected in xenografts *in vivo*

To identify potential differences in the transcriptome of leukemia cells in mono- and in co-culture, we preformed scRNA-seq of 6 different ALL PDXs under both conditions. In order to capture potential subtype heterogeneity, 4 different ALL subtypes were selected including two TCF3::HLF positive B-ALLs, two TCF3::PBX1 B-ALLs, one cortical T-ALL and one pre-T-ALL. In addition, these PDXs were also extracted and sequenced directly from the murine bone marrow (referred to as *in vivo*) with the aim of identifying potential similarities between those conditions (Fig. 5A). All the cells were successfully integrated and clustered based on the ALL subtype (Fig. 5B, suppl.Fig.3A). Differential gene expression analysis (DGEA) revealed upregulation of genes that are associated to B- and T-cell development as well as leukemogenesis when comparing co-cultured to mono-cultured ALLs. The corresponding dot plots highlight the similar expression of these genes *in vivo* and in co-culture, suggesting that the *ex vivo* model recapitulates essential features for leukemic development (Fig. 5C,D).

**Figure 5.**
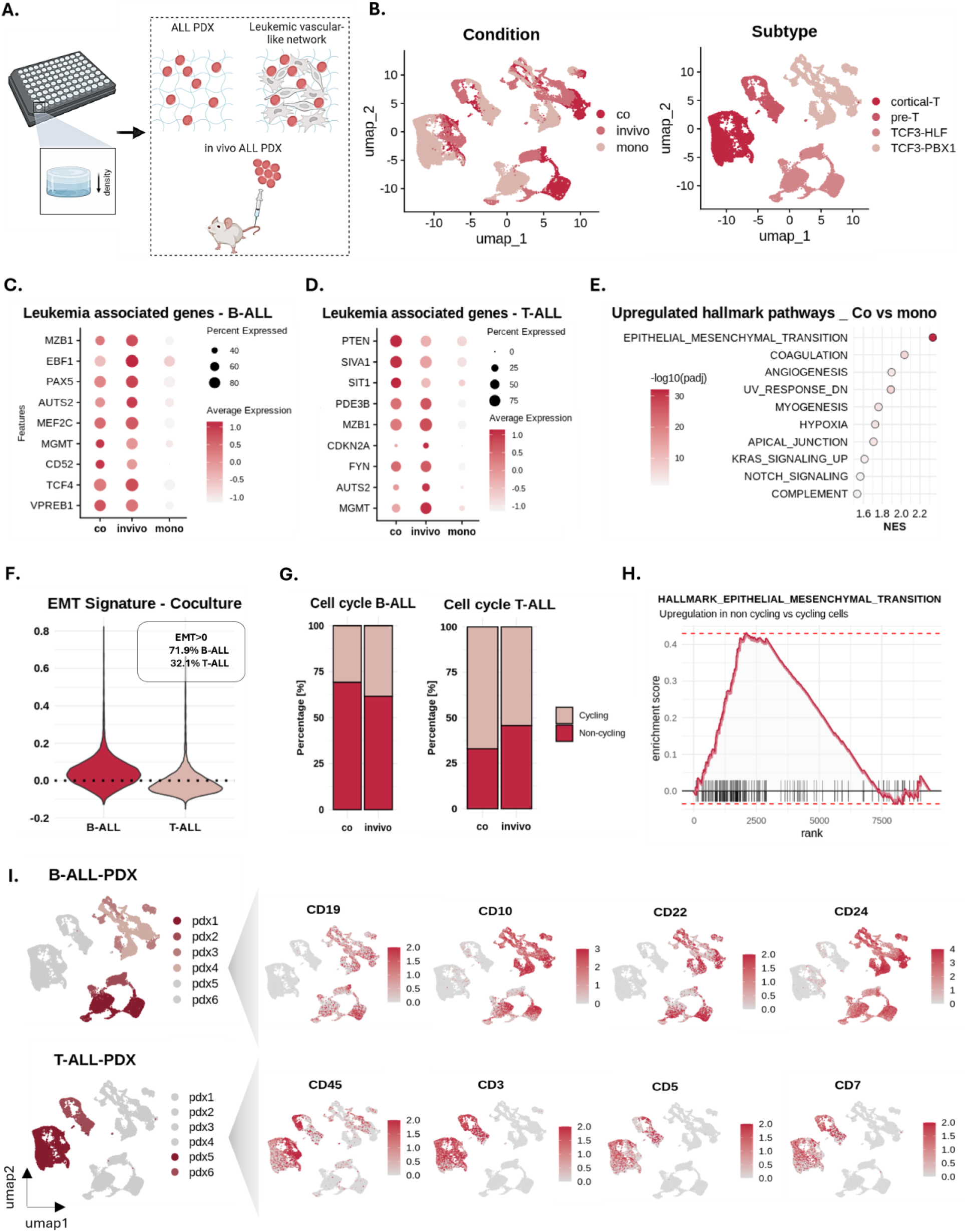
Comparison of leukemic gene expression from 3D co-culture and in vivo leukemia xenografts. A. Sequencing of leukemic PDXs in three different conditions, mono-, co-culture and bone marrow cells from leukemia xenografts in vivo. B. UMAP projections of the combined datasets based on the condition (left) and the subtype (right). The subtype is the leading source of variability, determining the clustering in the UMAP projection. C. Expression of leukemia-associated genes for B-ALL. D. Expression of leukemia-associated genes for T-ALL. E. GSEA comparing co-cultured ALL cells to mono-cultured. Depicted are the top 10 upregulated hallmark pathways. EMT is revealed as the most upregulated pathway in co-cultures. F. Violin plot showing the expression of the EMT signature in B- and T-ALL cells for the co-culture, with B-ALL cells having overall increased expression. G. Stacked bar plots showing percentages of cycling and non-cycling cells in B- and T-ALL for the two conditions, co-culture and in vivo. H. Enrichment plot of EMT comparing non-cycling (G1) to cycling cells (G2M and S). I. UMAP projections depicting the B-ALL (top, pdx1-pdx4) and T-ALL (bottom, pdx5-pdx6) PDXs, linked to expression of known ALL cell surface markers.

Further comparison revealed that genes regulating migration, adhesion and ECM-remodeling are amongst the most upregulated genes in co-cultured ALL (suppl.Fig.3B). The epithelial-mesenchymal transition (EMT) hallmark, described by some of the previously mentioned genes, was found to be the top pathway enriched, while the overall regulatory mechanism of EMT was consistently upregulated as well (Fig. 5E, suppl.Fig.3C). Notably, in recent studies the EMT pathway has been associated with increased aggregation and invasiveness in the context of leukemia, reminiscent of the enhanced motility upon immediate ALL- microenvironment interaction we have observed (Fig.2A-E).^44–48^ Independent analysis of B- and T-ALL cells validated EMT enrichment in both co-cultured subtypes, while this signature was predominant in B-ALL with approximately 70% overall expression (Fig. 5F, supp.Fig.3D,E). B-ALL has been described by enhanced microenvironment interactions *in vivo* compared to T-ALL, in line with its dependency on an EMT-like phenotype.^10^ Moreover, association of cell cycle arrest and EMT-like leukemic populations has also been described.^47^ Cell cycle analysis revealed that the 3D co-cultures successfully reiterate *in vivo* subtype-based proliferation heterogeneity with similar proportions of cycling and non-cycling cells; however, GSEA revealed that the EMT pathway is actually predominantly regulated by the non-cycling cells, potentially confirming a connection between cell cycle arrest and the EMT leukemic traits (Fig. 5G,H, suppl.Fig.3F,G). Finally, subpopulation heterogeneity based on known cell surface markers was detected, with cells exhibiting various expression profiles across the same PDX (Fig. 5I).

### Preserved cell heterogeneity in co-culture is validated by single cell microscopy and flow cytometry

In order to investigate whether an EMT-like, i.e. aggregated and migratory, phenotype is indeed present in the co-cultures, phenotypic and functional characterization was performed. For that purpose, thirteen PDXs of different subtypes including TCF3::HLF, TCF3::PBX1, B-other and T-ALL were seeded in both mono- and co-culture and analyzed using confocal microscopy and flow cytometry after 72 hours (Fig.6A). Single-cell distance analysis revealed increased aggregation in the co-culture, with the majority of the leukemic cells maintaining close distances of approximately 10μm (Fig. 6B, suppl.Fig.4A). Subtype-based analysis revealed enhanced clustering in B-ALL compared to T-ALL, associating B-ALL to the observed EMT increase in our transcriptomic data (Fig. 6C, Fig. 5F). In addition, migration of the ALL cells in the hydrogel was promoted by the non-hematopoietic cells when compared to mono-cultures, where minimum migration is detected and only a subset of T-ALL cells exhibited enhanced migratory capacity, yet again confirming T-ALL versatility (Fig. 6D,E). On the other hand, migration is observed in the case of all co-cultures (Fig. 6F). The portion of leukemic cells, in direct contact with the vasculature-like network, was assessed based on the HUVECs migration capability, with the network settling at a depth of 40% distance from the top (with the highest leukemia cell as position reference) regardless of the ALL subtype used (Fig. 6G). No leukemia cells were found in this depth in mono-cultures, while 20-40% of the co-cultured ALLs were able to migrate closely to the network, irrespective of the subtype (Fig. 6H, suppl.Fig.4B,C).

**Figure 6.**
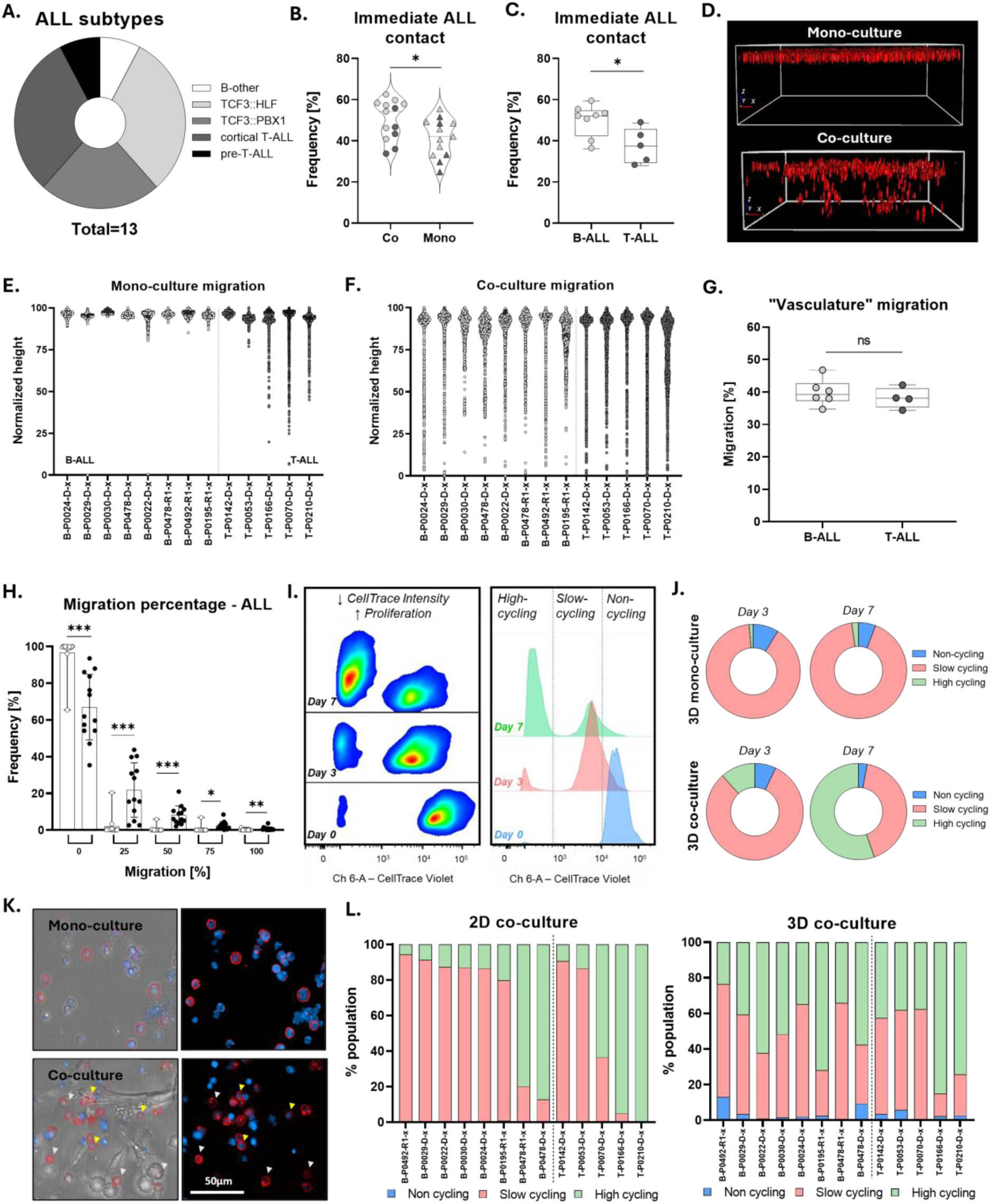
**3D co-culture-enhanced ALL aggregation, migration and proliferation lead to increased heterogeneity.** A. 13 PDXs of different subtypes were analyzed in the 3D model. B. Immediate contact, in less than 10μm, is enriched in the co-cultures, suggesting superior cell communication (statistics: Wilcoxon matched-pairs signed rank test, p value <0.05, triplicates for all conditions, PDXs used as biological replicates). C. Immediate contact is also enriched in B-ALL samples compared to T-ALL (same statistics as B). D. 3D reconstruction of B-ALL cells in the hydrogel. E,F. Single-cell distance analysis was performed, normalization to the highest ALL z-position in each well. Light gray: B-ALL PDXs, Dark gray: T-ALL PDXs. G. Dot plot showing the migration percentage of the HUVECs, normalization to the highest leukemic cell detected in the hydrogel (statistics: unpaired t test, p value <0.05, triplicates for all conditions). H. Migration analysis reveals that approximately 20-40% of ALL cells are in direct contact with the vessel-like structure (statistics: Wilcoxon matched-pairs signed rank test, p value <0.05, triplicates for all conditions). The migration percentage was calculated based on the cell position and the highest ALL cell position detected. I. Proliferation assessment with CellTrace Violet via Flow Cytometry. The CellTrace intensity reveals a non-cycling (defined by timepoint 0, high CellTrace intensity), a slow cycling (medium CellTrace intensity) and a high cycling population (low CellTrace intensity). J. Plots depicting the percentage of cells in each phase combined for all 13 PDXs per condition. K. Immunofluorescent staining with CD19 for lymphocyte (B-ALL) identification. Both CellTrace positive (yellow arrows, non-cycling) and negative cells (white arrows, cycling) are detected. L. Stacked bar plots depicting the percentage of non-cycling, slow cycling and high cycling cells in 3D and 2D co-culture respectively.

Furthermore, the leukemic PDXs were stained with CellTrace Violet and their proliferation was monitored for 7 days (Fig. 6I, suppl.Fig.4D). On day 3, no significant difference is observed across the conditions or subtypes, as expected based on the transcriptomic data (Fig. 6J, suppl.Fig.4E-G). However, at a later timepoint (day 7), increased proliferation is observed in the co-culture, showing beneficial conditions for leukemia’s growth. Interestingly, a portion of the leukemic cells decrease or pause their cellular growth rate, validating the existence of a non-cycling population as well as proving that the 3D co-culture maintains the cell cycle heterogeneity that is observed in *in vivo* studies (Fig. 6J). These findings were also validated by immunofluorescence, where upon CD19 staining for the lymphocyte identification, the presence of both CellTrace positive (yellow arrows, non-cycling) and negative cells (white arrows, cycling) was evident (Fig. 6K). Nonetheless, no specific localization based on the cycling state was observed, similarly to our previous *in vivo* observations where, upon chemotherapy, no selective cell death occurs based on the proliferation phase of the cells (suppl.Fig.4H-K).^10^ Finally, comparison of the proliferation rate of the ALL PDXs to the standard 2D hTERT MSC-ALL co-culture system revealed preserved cell state heterogeneity in 3D. Leukemic cells present a distinct proliferation phenotype in 2D with patient samples either strongly proliferating or completely lacking division capacity, while in 3D both cycling and non- cycling populations are detected for all PDXs, thus highlighting that the 3D model can recapitulate cellular complexity *ex vivo* when 2D systems fail (Fig. 6L).

### Drug response patterns are reproduced, with increased resistance in 3D

Finally, we investigated whether the 3D co-cultures can be utilized for drug response evaluations and if the distinct phenotypes observed alter drug sensitivity. Flow cytometry was performed to evaluate the 3D response. For that purpose, 16 PDXs of various subtypes were treated for 72 hours, with compounds of interest specifically for each subgroup, and then enzymatically extracted for evaluation. Flow cytometry analysis was performed by gating for CellTrace positive cells, i.e. lymphocytes, based on unstained controls, followed by propidium iodide gating for the exclusion of dead cells (Fig. 7A). The viability of the ALL cells was compared to 2D controls that were treated in the same manner and analyzed by high throughput microscopy (Fig.7B).

**Figure 7.**
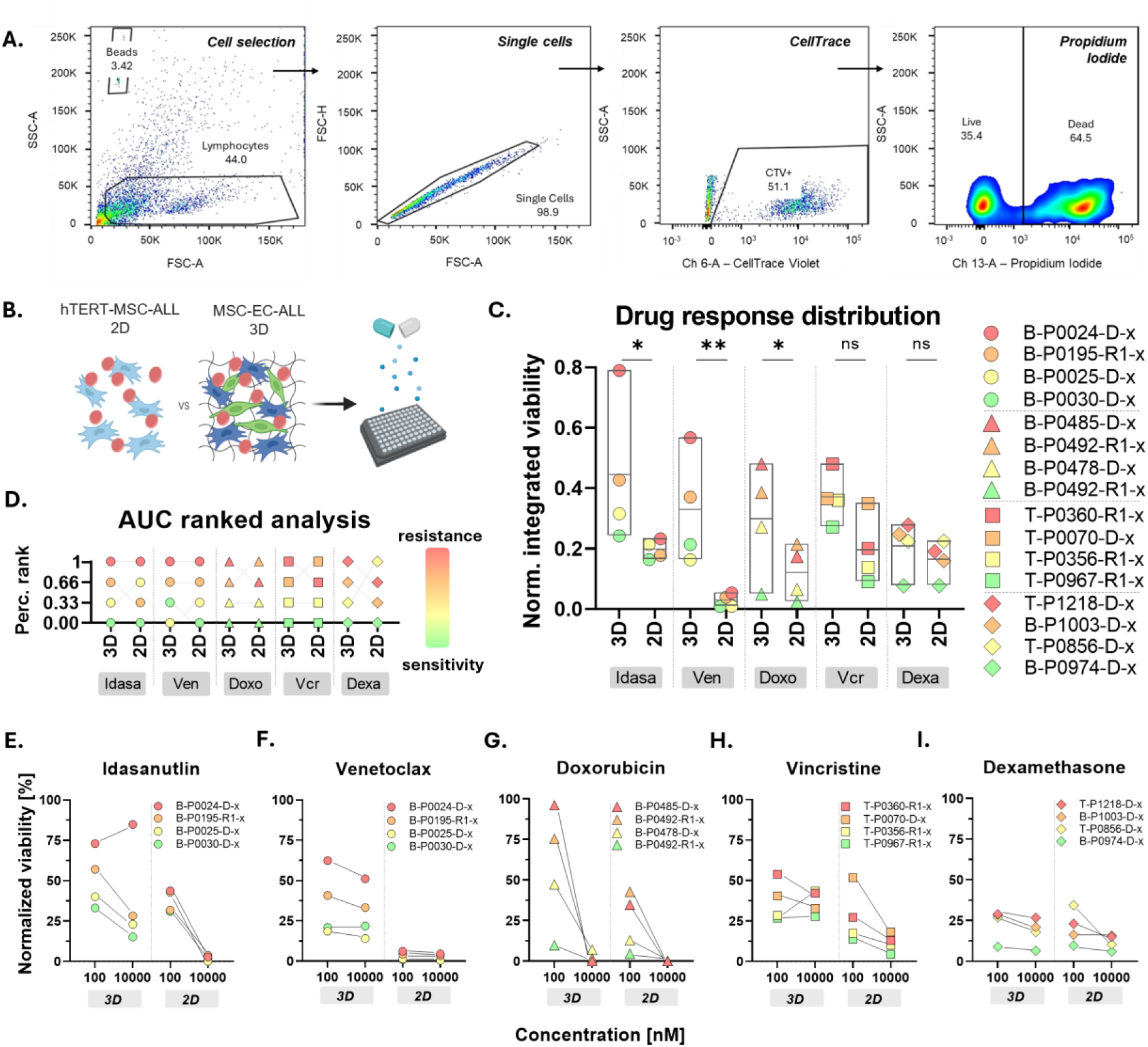
Drug screenings in the 3D co-culture reveal increased resistance. A. Gating strategy for flow cytometry. The living ALL cells were gaited for lymphocytes based on their size, followed by doublet exclusion, cellTrace positivity and PI negativity. B. Experimental design. The 3D co-culture response was compared to the standardized drug response profiling protocol, where immortalized hTERT-MSCs-ALL co-cultures are treated for 72hrs, in triplicates. C. Bar plots showing the normalized response values (integrated normalized cell counts over two concentrations) for all the 3D and 2D conditions (statistics, ratio paired t test, significant p value <0.05, triplicates). D. Percentile ranking indicating the response patterns across the conditions, color-coded based on the response in 3D. E-I. Dot plots showing the normalized viability of the leukemic cells upon treatment with different compounds. Depicted is the response to 100 and 10,000nM in the 2D and 3D systems respectively. From left to right, idasanutlin, venetoclax, doxorubicin, vincristine and dexamethasone plots.

The predictive value of the 3D model was validated as the samples were following the response patterns of the established 2D drug response profiling (DRP) platform (Fig.7C). Percentile ranking was performed to confirm the sensitive/resistant samples of each cohort (Fig.7D). Remarkably, there was increased viability in 3D upon treatment for the majority of the PDXs (Fig.7E-I). 14 out of 20 samples (70%) showed decreased sensitivity (>10% difference in normalized cell counts to DMSO) at 100nM for the 3D co- cultures, suggesting superior ALL protection upon treatment. This phenotype can be attributed to multiple factors, as the presence of both ECM and ECs have been linked to leukemic treatment escape.^26,49,50^ In addition, a remaining subpopulation can be observed at 10μM for 4 drugs (exception doxorubicin) indicating the existence of a resistant subpopulation that cannot be detected in 2D (Fig. 7E-I). Taken together, this model recapitulates established drug response patterns thus can be used for compound testing applications, while the decreased drug efficacy observed could also potentially lead to the identification of resistance patterns more readily compared to 2D.

## Discussion

Three-dimensional bone marrow models have demonstrated significant improvements in the study of leukemia, as they incorporate essential niche elements including various cellular types, ECM communication and mechanical support; however, they mostly focus on the study of acute myeloid leukemia (AML) and not ALL.^51^ In this study, a 3D hydrogel-based BM mimic for ALL PDXs is presented, characterized by robustness, reproducibility and compliance with high-throughput readouts. The model recapitulates blood-vessel like structures by culturing human primary MSCs and ECs with the ALL PDXs in a responsive hydrogel that facilitates cell-cell and cell-ECM communication. Subtype-dependent localization towards the vasculature-like structures was observed, with B-ALL requiring closer contact with stromal or endothelial cells whilst T-ALL does not preferentially localize close to the structures. Barz et al. demonstrated similar phenotypes in an *in vivo* xenograft model with B-ALL cells residing in proximity to vascular and endosteal niches in the BM, while T-ALL cells exhibited increased versatility.^10^ This highlights the capability of our *ex vivo* model to reproduce subtype-relevant interaction features observed *in vivo*.

The BM microenvironment impact on leukemic progression is well-described; however, whether the niche acts as a predisposition factor for disease or it is transformed into a favorable environment upon crosstalk with malignant cells is yet unclear.^52^ Single-cell transcriptomic profiling of the 3D cultures revealed multi- lineage differentiation of the MSCs in the vasculature-like system, with a polarization based on the CXCL12 and PDGFRA markers, linking the *ex vivo* phenotype to the ones described in human or murine models with distinct lineage commitment.^35–38^ Smooth muscle cells and fibroblasts were defined as the predominant lineages in the HUVEC-containing conditions, potentially due to their capability of producing vascular-promoting and supporting factors.^39,40^ No significant ALL-induced alterations were observed suggesting that longer culturing is necessary to monitor their crosstalk dynamics. On the other hand, HUVECs enriched the ECM degradation pathway upon leukemic influence, suggesting ECM re-organization to facilitate malignancy, as shown in Verma et al. that described how ECM degradation through the microenvironment influences leukemic progression.^53^ Thus, this *ex vivo* vasculature-like model can also be utilized to determine trajectory fates of supporting cells, as well as identify key mediators of their interdependence with leukemia.

The relevance of the co-culture system to leukemic *in vivo* features was demonstrated by comparable expression of necessary B- and T-cell developmental genes to PDX cells isolated directly from the murine BM. Notably, these genes were not significantly expressed when ALL cells were cultured alone in the hydrogel, highlighting the necessity of microenvironment elements for a leukemic 3D model, as shown in other studies.^51^ Furthermore, through further analysis of the leukemic profile in 3D co-culture, the EMT pathway was identified as the highest enriched. EMT-like leukemia has been described by increased invasiveness in multiple studies, with Park et. al recently showing that MSC-adherent ALL exhibits enhanced survival and resistance to treatment compared to non-adherent cells through EMT-related genes overexpression.^44–48^ Single-cell topological analysis revealed enhanced aggregation and migration in the 3D co-culture. Interestingly, only a distinct subpopulation (20-40% of ALL cells) exhibited these properties thus highlighting preserved heterogeneity in the BM mimic. Subtype-based variability was detected with B-ALL exhibiting increased dependency on the niche, with enhanced aggregation in co-culture as well as limited migration capability when cultured alone compared to T-ALL. Cell cycle arrest has also been associated to the EMT-like phenotype and chemoresistance.^47^ Proliferation analysis revealed high or slow cycling cells, preserving cell state heterogeneity that our 2D model was incapable of. Taken together, the 3D BM mimic has the capacity of recapitulating transcriptomic signatures and exhibiting subpopulation heterogeneity similar to the one observed in the BM.

Two-dimensional drug screening applications are widely used in cancer research; however, there is still a gap in translational medicine as only 5% of the FDA approved compounds show any efficacy in clinical trials.^54^ 3D systems have thus been implemented as they better recapitulate *in vivo* complexity.^55^ Here, we examined the predictive value of our system by comparison to the established Drug Response Profiling (DRP) platform, since it has been implemented for informed personalized therapies and identification of recurrent sensitivity patterns for subsets of ALL PDXs for personalized treatment options.^56–58^ The drug response successfully emulated the PDX distribution (sensitivity/resistance), which highlights the relevance of our 3D drug testing pipeline. Increased resistance was observed in 3D for all compounds, with PDX-dependent variability. Significantly, at 10μΜ, resistant leukemic subpopulations were identified for 4 out of 5 drugs tested. This is in line with numerous reports that showcase that the enhanced complexity of leukemic 3D models leads to increased resistance that potentially mimics better the clinical response.^13,15,59–61^ Increased resistance could be attributed to the altered drug distribution in 3D and to the presence of both the ECM-mimic and the ECs, potentially suggesting a more representative system for drug distribution compared to 2D.^62^

Further investigation is needed to decipher the molecular drivers of this resistance and to identify distinct leukemic subpopulations. scRNA-seq upon treatment could be implemented to unravel the *ex vivo* complexity, while comparison with *in vivo* treated PDXs (with our established murine PDX model) could identify minimal residual disease patterns and mechanisms. We have already shown that *in vivo* disease signatures can be recapitulated in our *ex vivo* model. If signature similarities are maintained upon treatment as well, we will potentially have a tool to assess treatment options and study resistance patterns without the use of murine models. Furthermore, the topological and phenotypical characterization of the ALL PDXs is needed to complement the flow cytometry drug readout. Identifying resistance patterns related to the position, proximity or phenotypic alterations of the cells will lead to novel treatment options by interfering with the leukemia-microenvironment axis. It will also facilitate the use of this system for complex drug combinations screenings and functional genetic studies.

Taken together, we have established a 3D vasculature-like model that reproducibly mimics physiological-like features. Specifically, our model identifies distinct leukemic subpopulations that are characterized by different proximity to the supporting cells, diverse stiffness selection and even multiple proliferation states. These features cannot be detected in 2D systems as cellular heterogeneity is completely lost. We have also shown that drug response can be studied in this system, systematically recapitulating 2D response patterns for each PDX, with increased resistance, thus potentially allowing the identification of relapse-driving populations. These systems can bridge the gap between *in vivo* and *ex vivo* and can potentially lead to profound understanding of different mechanisms related to disease progression, microenvironment interactions and therapy resistance.

## Materials and Methods

### Patient-derived xenografts

The primary patient samples were collected through bone marrow aspiration upon written informed consent of the patients’ parents or legal guardians, in accordance with the Declaration of Helsinki. The ethics commission of the Canton of Zürich granted the approval for the sample collection with the approval number 2014-0383. The primary samples were then injected (fresh or frozen) via intravenous injection in immunodeficient NSG (NOD.Cg-*Prkdcscid IL2rgtm1Wjl*/SzJ) mice, that were purchased from The Jackson Laboratory. The human leukemic engraftment was monitored weekly by flow cytometry and xenografts were recovered by samples enrolled in the ALL-BFM 2000 and 2009 and ALL-REZ BFM 2002 studies.

### Cell culturing

The primary human mesenchymal stromal cells (MSC) were provided by Prof. Dr. Martin Ehrbar. The extraction of the MSCs through bone marrow aspiration of healthy donors was previously described in Papadimitropoulos et al, 2014.^63^ The cells were cultured in MEMa (22571-020, ThermoFisher) supplemented with 10% FBS (S0615, Sigma), 1%PS (15140122, ThermoFisher) and 5ng/mL FGF-2 (PHG0369, ThermoFisher) at 37°C with 5% CO_2_.

The Human Umbilical Vein Endothelial Cells were also provided by Prof. Dr. Martin Ehrbar, purchased from Angioproteomie (PELO Biotech). The cells were cultured in EGM-2 (CC-3162, Lonza) supplemented with 10% FBS in collagen-coated (A1048301, ThermoFisher) flasks at 37°C with 5% CO_2_. Both unlabeled (PB-CH-190-8011) and GFP-transduced (PB-CAP-0001GFP) HUVECs were used for the formation of vascular-like networks.

### 3D culturing

The 3D co-culture was established in the 3DProSeed 96-well hydrogel plate (ECT.PS1.001.096, Ectica Technologies), a black, glass-bottom imaging plate with 96 wells containing pre-cast, synthetic, animal-free, and optically transparent PEG-based hydrogels. Cell suspensions were prepared in MEMα supplemented with 10% FBS, 1% PS, and 5 ng/mL FGF-2. The hydrogel storage buffer was aspirated from the plate and discarded, and 200 μL of cell suspension was added to each well, followed by incubation at 37°C with 5% CO₂. For the MSC-HUVEC-ALL co-culture, 30,000 primary human MSCs were seeded per well initially. After 24 hours, 30,000 HUVECs, 30,000 ALL cells, and 15,000 MSCs were added to each respective well. MSC-HUVEC co-cultures (MSC were incubated for 24 hours, followed by seeding of HUVEC and MSC) and ALL monocultures served as controls, under the same experimental conditions. Vascular-like network formation required 72 hours following the addition of HUVECs. For long-term cultures (7 days), an additional 100 μL of medium was added on the third day.

### Hydrogel digestion

Cells were seeded as described above (3D culturing). At the digestion timepoint, the supernatant of each well was collected and 200uL of digestion mix was added. The mix included Liberase 0.2mg/mL (5401119001, Sigma) DNase 200U/mL (11284932001, Sigma) in MEMa. The mix was incubated for 30min at 37°C with 5% CO_2_, with a manual mixing step at 15min. The cell suspension was collected and 200uL of MEMa was added for the dilution of the digestion mix. The samples were centrifuged at 300g for 5 minutes and resuspended in the appropriate solvent.

### Cell staining

CellTrace agents i.e. CellTrace Violet, CellTrace Yellow and CellTrace FarRed (C34571, C34573, C34572, ThermoFisher) were used for the staining of the different cellular types for sequential microscopy analyses. The staining of the cells followed the manufacturer’s instructions, whereas the cells were stained for 20min in PBS at RT and washed for 5min, followed by resuspension in medium and seeding.

### Immunostaining

Anti-human CD90 conjugated to FITC or APC (11-0909-42, ThermoFisher) were used for the staining of the MSCs. Anti-human CD31 conjugated to FITC or Pacific Blue (557508 BD biosciences, 303114 Biolegend) were used for the HUVECs. 1:300 of antibody was directly added in the medium of the well and incubated for 30min. For the leukemic cells, antibodies were selected based on the subtype, with CD19 APC (345791, BD Bioscience) for B-ALL and CD7 PE (12-0079-42, Thermofisher) for T-ALL. 1:600 of antibody was directly added in the medium of the well and incubated for 2hrs and 30min.

### Confocal microscopy

High-content confocal microscopy was performed with an Operetta CLS (PerkinElmer). The objectives used were 10x, 20x and 40x with z-stack acquisition. The standard step size for 10x imaging was 7.8um. All conditions were imaged in triplicates. Time-lapse imaging was performed with 10x magnification, 5 minute time interval and 10um step size for the z-stack acquisition.

### Confocal microscopy analysis

All microscopy analysis was performed with Harmony (version 4.9), the Operetta CLS software. All of the images were loaded either as a 3D reconstruction or a maximum projection (based on the requirement of the downstream analysis) and each cellular type was segmented with the appropriate algorithm model. For the calculation of 3D volume and surface, the *find image region* function is implemented and the segmentation mask is created based on the pixel intensity in relation to the local or absolute threshold. The choice of the threshold depends on the fluorophore used. For the calculation of the position in z, the *find nuclei* function is performed for the 3D segmentation of leukemic cells, while for the other cell types *find image* region is selected as above, and the *calculate position properties* building block results in the quantification of the centroid’s position of each single cell/region in the well. Following the same principle, the cross population properties (minimum distance to nearest neighbor) are identified by calculating the distance from the selected object’s border to the object border of the nearest neighbor object. Finally, 2D motility analysis is performed through the *calculate kinetics properties* building block, with current step size calculated by (S_-1_ + S_+1_) and current speed by (s_-1_ + s_+1_) / (t_+1_ + t_-1_), with s as the position and t as the corresponding time point.

### Proliferation analysis

ALL cells were stained with CellTrace Violet and the seeding was performed as described above (3D culturing). After 3 and 7 days, the hydrogels were digested and the cell suspensions were resuspended in 100uL PBS each. 1uL of 7-AAD was added in each sample and left for 5min, for further exclusion of dead cells. Flow Cytometry experiments were conducted at a BD LSRFortessa (BD Biosciences) and all flow cytometry data were analysis using the FlowJo software (version 10.8.1).

### *In vivo* experiments

Tail intravenous injection of ALL patient-derived xenografts into unconditioned 6 to 8-week-old mice according to animal care regulations after approval by legal authorities (125/2013,124/16, 131/19) was performed for the PDX expansion. The level of human leukemia was monitored weekly by peripheral blood sampling. The blood samples were stained with anti-human CD45 Alexa Fluor 647, CD19 PE or CD7 PE(304018 Biolegend, 302208 Biolegend, 12-0079-42 Thermofisher) and anti-mouse CD45 eFluor450 (48- 0451-82, Invitrogen) with 1:100 dilution and analyzed with flow cytometry for the engraftment calculation.

### Sample preparation for single-cell RNA sequencing

For the 3D samples, cells were seeded as described above (3D culturing). After 72hrs, the hydrogels were digested and cell suspensions were collected. For the sample preparation, the protocol for the fixation of cells for chromium fixed RNA profiling (Chromium Next GEM Single Cell Fixed RNA Sample Preparation Kit, 1000414, 10x Genomics) was followed based on the manufacturer’s instructions. Briefly, the samples were fixed for 24hrs at 4°C, counted with an automated cell counter EVE^TM^ Plus (NanoEntek) with the use of trypan blue and resuspended in glycerol (G5516, Sigma) for long term storage.

For the *in vivo* samples, the femurs were harvested and the leukemic cells were flushed through the bone marrow and cryopreserved. Upon thawing, the samples were sorted using a BD FACS Aria to ensure leukemic population purity, with the use of CD19 PE (302208, Biolegend) for B-ALL and CD7 PE (12-0079- 42, Thermofisher) for T-ALL immunostaining. Samples were stained with trypan blue and counted manually with a hematocytometer. Finally, samples were resuspended in PBS containing 0.04% BSA.

### Library construction for single-cell RNA sequencing

Samples were multiplexed (1000456 for *ex vivo* samples and 1000261 for *in vivo* samples, 10x Genomics) and pooled to a total of 10000 cells per run. Then, they were loaded and sequenced with the use of an Illumina platform. The standard library protocol for 10x genomics single cell RNA sequencing for fixed and fresh cells respectively was followed based on the manufacturer’s instructions. 59,931 cells (35,545 leukemic) were recovered for the *ex vivo* samples, while 7,707 leukemic cells for the *in vivo* and were further processed computationally.

### Single-cell RNA data analysis

The processing of the raw sequence data was performed with CellRanger, using the reference genome library hg19.The data were analyzed with the R software (version 4.1.2) according to the standard pipeline described in the Seurat package (version 5.0.3). The pre-processing includes filtering of low-quality cells, doublet exclusion using the DoubletFinder package (version 2.0.3) and data integration for the *ex vivo* and *in vivo* datasets. Further analysis included gene set enrichment analysis (GSEA package, version), cell cycle scoring and identification of inter-cellular communication patterns (CellChat package, version 2.1.2). All p values > 0.05 were considered significant. Further information can be found in the published code (code availability).

## Code availability

The code used for the analysis can be found on GitHub: https://github.com/mkanari/scRNAseq_analysis

## Drug treatment

The compounds vincristine, doxorubicin, dexamethasone, venetoclax and idasanutlin (S1241, S1208, S1322, S8048, S7205; Selleckchem) were directly added to the well supernatant with the use of the D300 drug dispenser (Tecan) after 72hrs of culture. The concentrations used were 100, 1000 and 10000nM, depending on the experimental conditions. The drugs were incubated for 72hrs, followed by a microscopy (Operetta CLS, PerkinElmer) or flow cytometry (BD LSRFortessa, BD Biosciences) readout.

The flow cytometry readout included digestion of the hydrogels as described above. The samples were resuspended in 100uL PBS. 1uL propidium iodide (P1304MP, ThermoFisher) was used per sample for 5min to determine the dead cells. For the viability quantification, 5uL AccuCount fluorescent counting beads (ACFP-70-10, Spherotech) were added per sample. All conditions were normalized to DMSO controls (10uM, D8418, Sigma).

## Two-dimensional drug response profiling

Two-dimensional drug response profiling followed the established protocol from DRP Zurich, described in Frismantas et al., 2017.^56^ 2500 h-TERT MSCs in AIMV (C35012, ThermoFisher) were plated in a 384 well plate. After 24hrs, 10000 leukemic cells in AIMV were seeded on top of the MSCs for the establishment of co-cultures. The following day, the drug compounds were added using the D300 drug dispenser (Tecan) at the same concentrations as above for 72hrs. For the viability determination, CyQuant staining (CyQUANT Cell Proliferation Assay, C35012, ThermoFisher) was performed for 45min (1:40 dilution for suppressor, 1:600 dilution for compound, MEMa).

## Statistics

All statistical analyses were performed using the software GraphPad Prism (Version 10.0.2). P values < 0.05 were considered significant. The statistical tests performed were chosen based on each data set and are indicated at the figure legend of the corresponding graph.

## Authors’ contributions

MK designed, conducted experiments and analyzed data. FDS and LAK provided support in data interpretation. MK and IJG performed sequencing experiments. CB, BD and ON contributed to the single- cell RNA sequencing experiments. MK analyzed the sequencing data. IJG performed the xenograft experiments. BRS assisted in the development of the 3D cell culture system. ME, BB and J-PB conceptualized the project. BB and J-PB supervised the study; MK, BB and J-PB wrote the manuscript. All authors approved the final version of the manuscript.

## Data sharing statement

The data supporting the findings of this study are available from the corresponding author upon request to J.-P.B. The single-cell RNA sequencing data will be made available from NCBI GEO upon publication (or request of the editor/reviewer).

## Supporting information

supplementary_material_Kanari

## Acknowledgement

The authors would like to acknowledge the Functional Genomics Center Zurich (FGCZ) of University of Zurich and ETH Zurich for performing the sequencing experiment. We would also like to thank the Center for Microscopy and Image Analysis, University of Zurich for supporting the image analysis. Sorting and sequencing of the *in vivo* samples was performed with the support of the Flow Cytometry Core Facility and the Gene Expression Core Facility at EPFL respectively. We would also like to thank Dr. Maria Mitsi for the fruitful discussions regarding the methods establishment. Finally, schematics were created with Biorender.

