## supplementary_material_Kanari for "A three-dimensional *ex vivo* model recapitulates *in vivo* features and unravels increased drug resistance in childhood acute lymphoblastic leukemia"

***Suppl. Fig.1: “Vascularization” occurs only in the case of coculturing MSC and EC cells and in the presence of primary stromal cells.***

*A. Maximum projections of 3D cocultures (z-step size 10μm), left depicting MSC, right HUVEC and ALL in both (MSC in blue, HUVEC in green, ALL in red). B. Maximum projection of 3D cultures. Leukemic cells (red) were cocultured with MSCs (blue, left) and HUVECs (green, right) separately. No network can be formed in either condition as the MSCs act as a scaffold for the network formation while no vascular-like network can be formed in the absence of endothelial cells. C. Immortalized hTERT-MSCs were cocultured with HUVECs and leukemic PDXs with the described protocol (3-D culturing). Upon 72 hours, maximum projections show no network formation, further reinforcing the observation that primary cells are needed to recapitulate physiological-like conditions.*

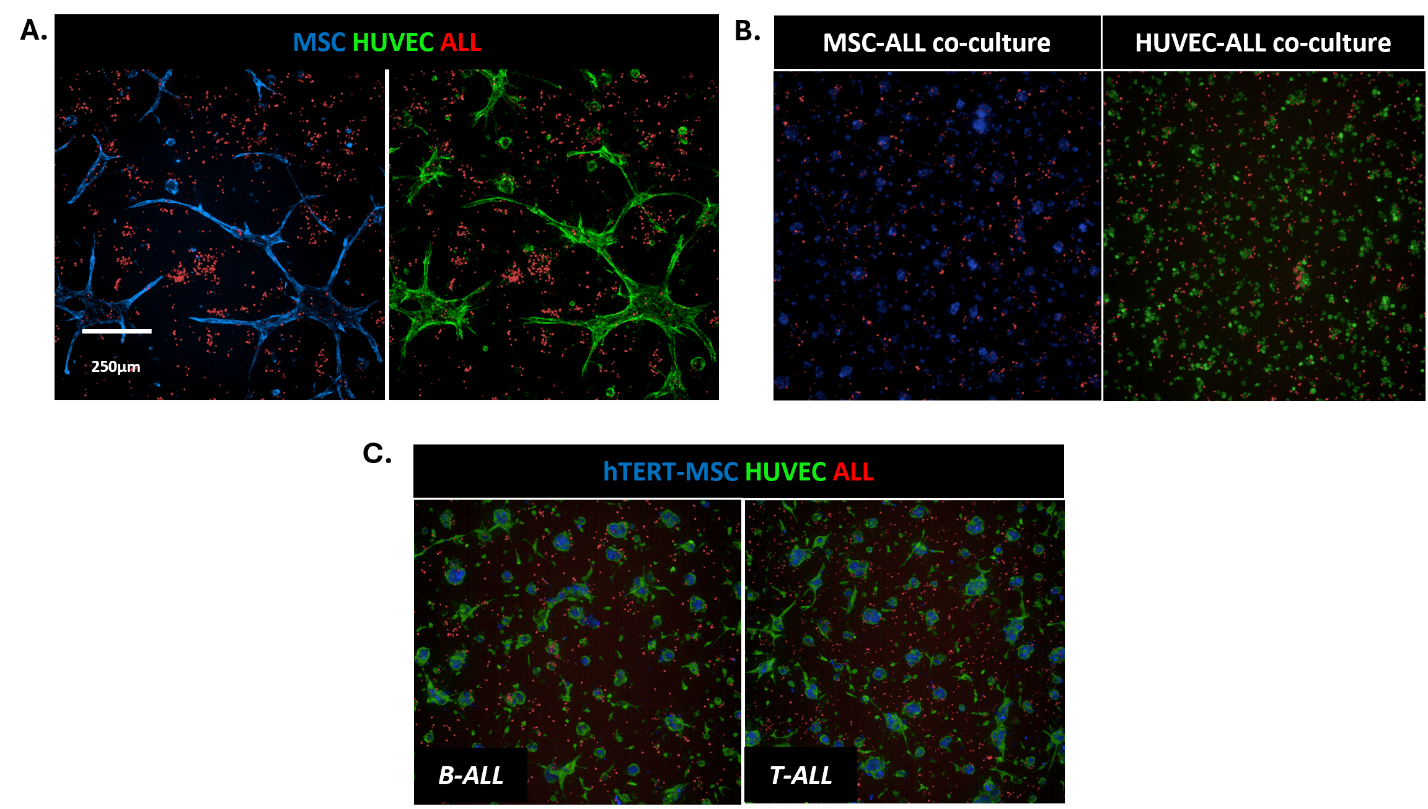

***Supp. Fig.2 suppl: Supplementary material for the supporting cells***

*A. MSC subpopulations. Dot plot showing the expression of known MSC markers that are used for the identification of distinct subpopulations. PDGFRA, PDGFRB, CXCL12 and LEPR expression is shown as they are considered known MSC markers with distinct expression. B. Stacked bar plot depicting the population distribution, as percentages, across the experimental conditions (MSC only, MSC vascular, MSC leukemic). C. Table showing the percentages of the MSC subpopulations across the conditions. D,E: GSEA comparing the conditions separately. Left: Vascular vs MSC only, right: leukemic vs MSC only. F: Analysis of cell-cell communication networks through the CellChat package. Bubble blot depicting the interaction axes between MSC and ALL, with the communication probability. Arrows indicating the most probable communications. G. GSEA comparing vascular vs HUVEC only. Pathways related to HUVEC replication are upregulated in the vascular condition, validating phenotypic observations as HUVEC display very low viability when cultured alone in the hydrogel.*

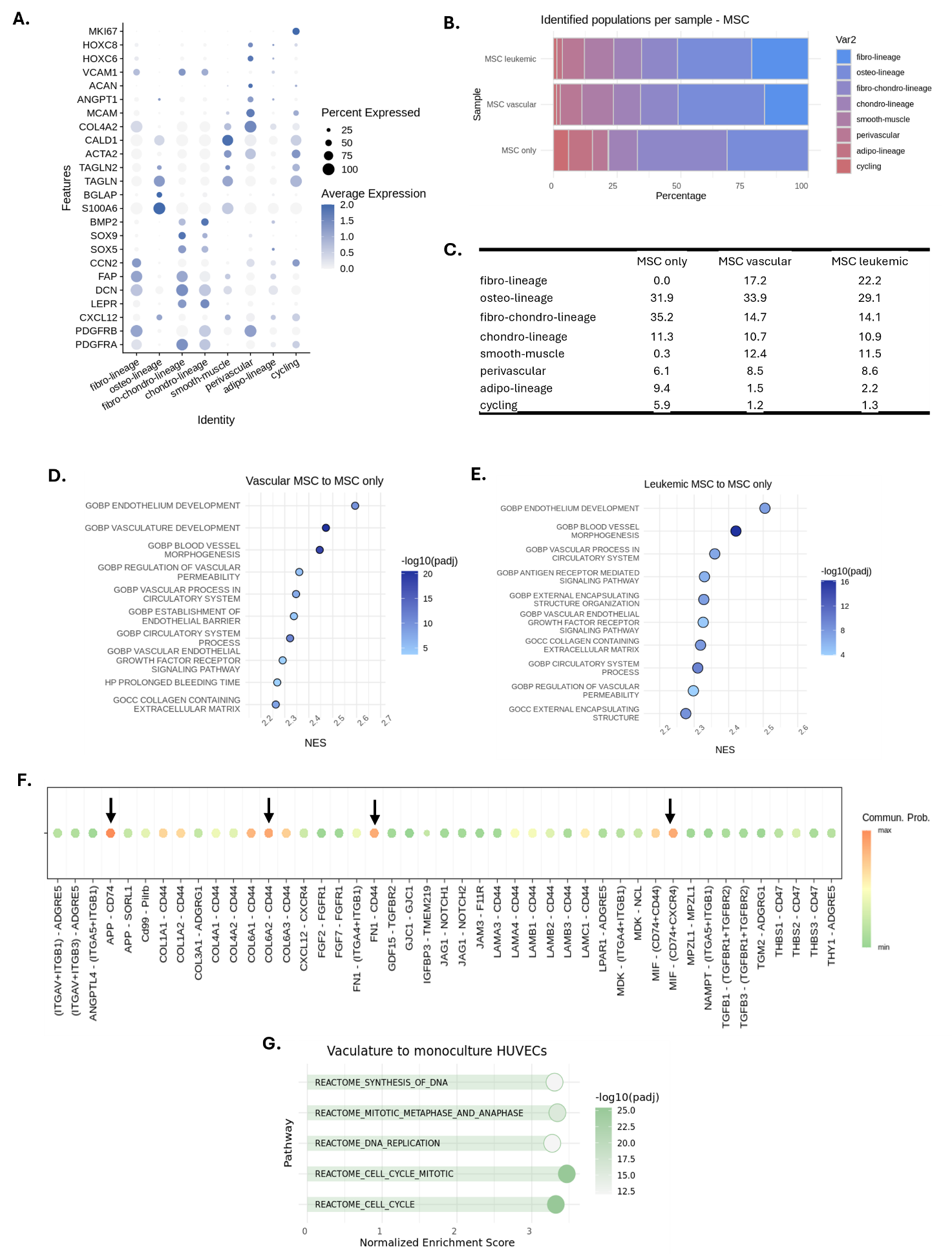

***Suppl. Fig.3: Supplementary material for leukemic PDXs.***

*A. UMAP projections of the ALL PDXs before (left) and after (right) integration. The conditions are initially separated based on the sequencing batch, with co- and mono-cultures clustering together. Upon integration based on the condition, the samples successfully integrate based on the leukemic subtype. B. Bubble plot showing the upregulation of EMT-related genes in co-cultures C. GSEA revealed that all the pathways the regulate the epithelial-mesenchymal transition pathway are upregulated in co-culture when comparing to mono. D. Subtype- based GSEAs showing that the EMT pathway is independently upregulated in co-culture for both B- and T-ALL. E. Cell cycle analysis plotted on the UMAP projection. F. The EMT signature was found to be upregulated in the non-cycling cells, which is in line with observations in the literature linking the EMT-like phenotype to quiescence.*

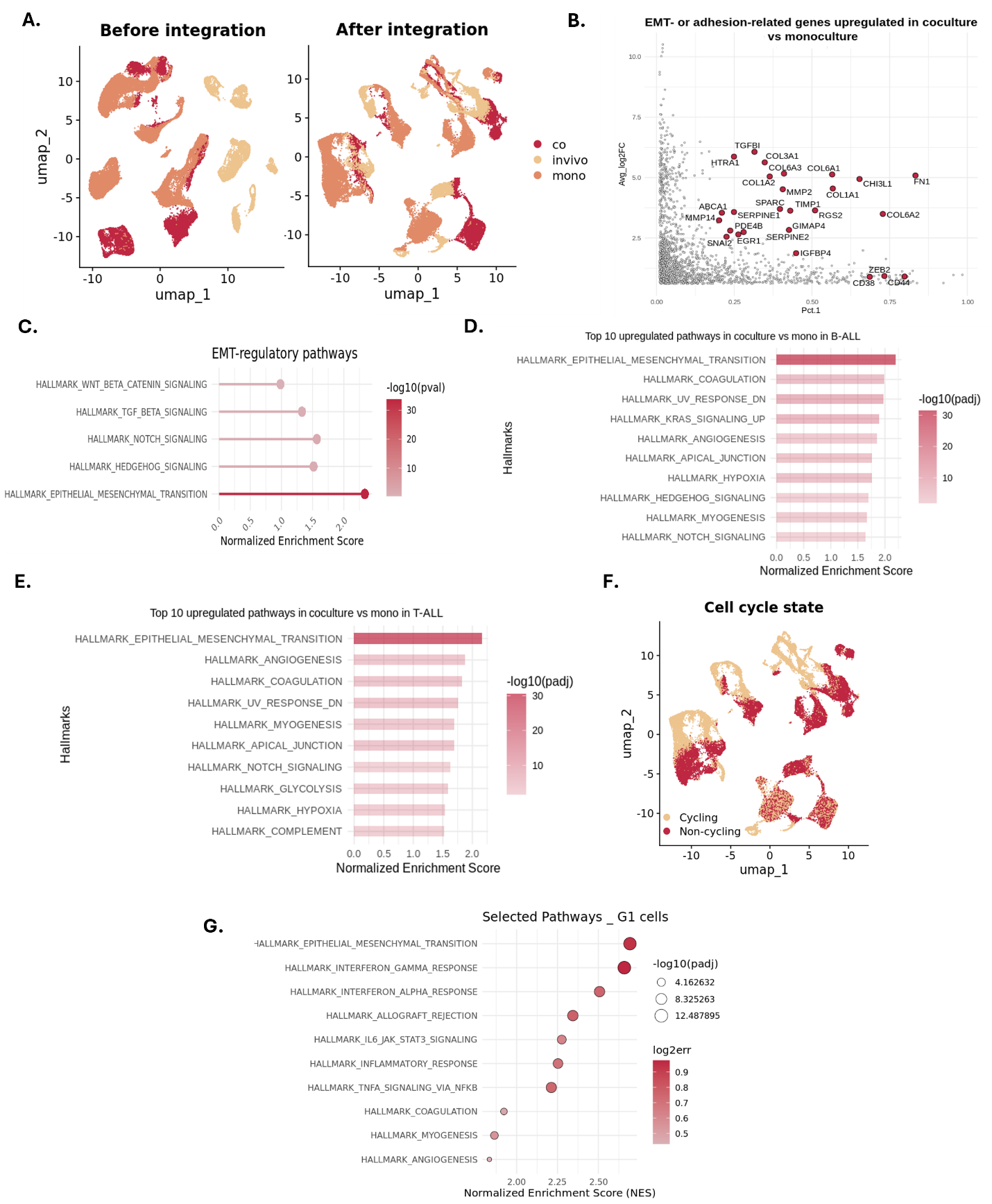

***Suppl. Fig4: ALL proliferation patterns and position in the hydrogel.***

*A. Histograms of ALL aggregation measured by single-cell analysis in mono- and co-culture. Data are represented by triplicates for each PDX with bin center = 4. B. Subtype-based migration percentages. Left: B-ALL PDX migration, right: T-ALL PDX migration. C. Subtype based migration percentages (bin center =10). Left: Mono-culture migration, right: Co-culture migration. D. Examples of CellTrace histograms for 4 leukemic PDXs. All of the PDXs exhibit increased proliferation capability on day 7; however, their kinetics differ based on the patient. E. Subtype proliferation comparison did not reveal significant differences across the two subtypes. Cells are classified as cycling (G2M/S phase) and non-cycling (G1) phase and the corresponding percentages are plotted as a stacked bar plot across the different conditions. No significant differences can be detected, showing that the cells proliferate in a similar manner ex vivo and in vivo. G. Stacked bar plots showing the proliferation percentage for (left to right): 3D mono-culture, day 3 and day 7, 3D co-culture day 3 and 2D co-culture day 3. H. Immunofluorescence images showing the top (left) and the bottom (right) of the hydrogel (blue MSCs, green HUVECs, red ALL). I. The migration percentage of each leukemic cell was calculated based on the highest ALL position. Cells that have migration capability higher than 25% are considered cells at the bottom. J,K: Histograms showing the distribution of CellTrace intensity on the top vs bottom cell (bin center = 0.1). No significant differences are observed, indicating that there is no preferential localization based on the cycling/non-cycling state.*

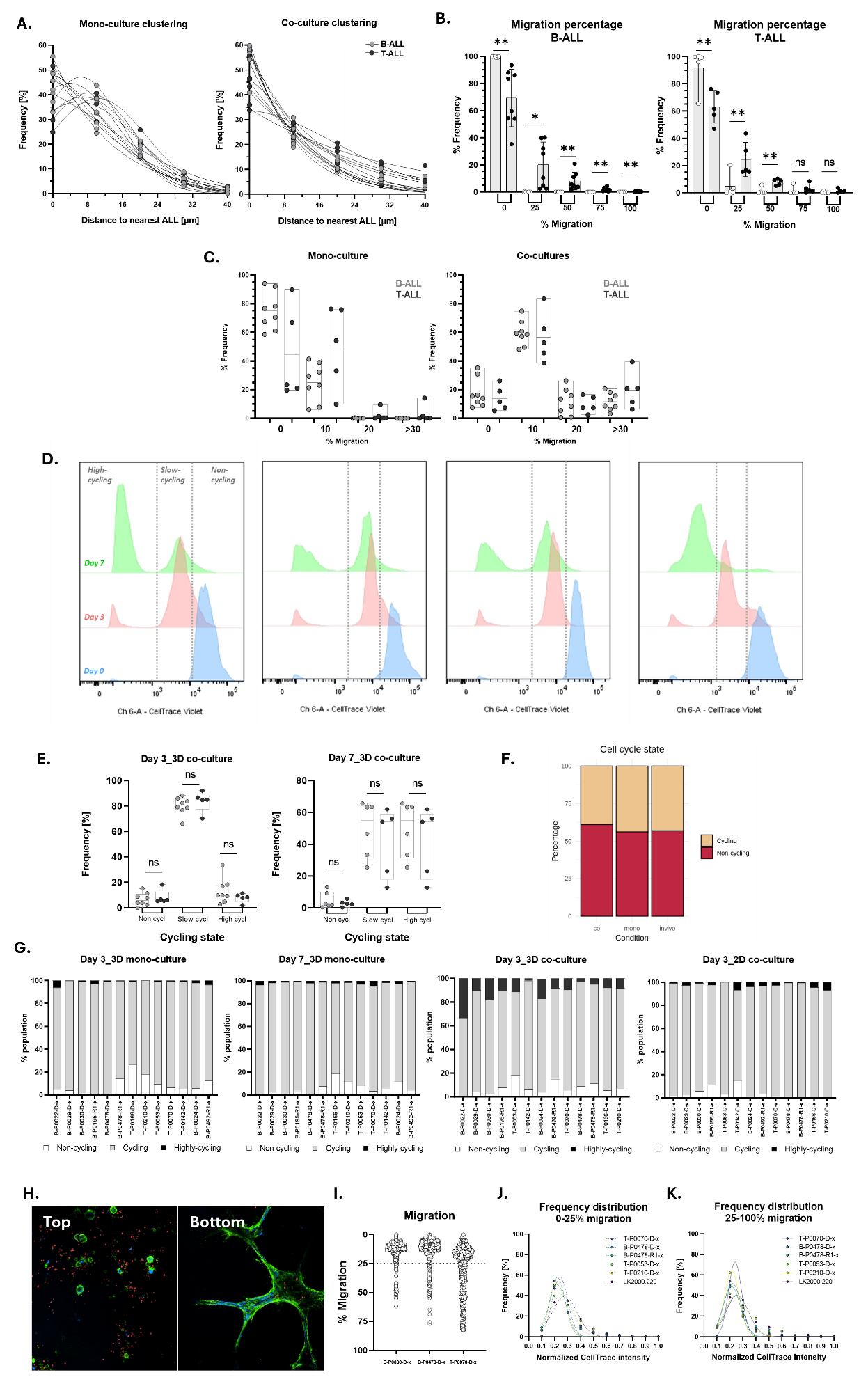

**Supplementary table 1: Patient-derived xenografts information**

| **PID** | **diagnosis** | **Disease stage** | **EGIL** | **subtype** | **genetics** |
| --- | --- | --- | --- | --- | --- |
| B-P0022D-x | BCP-ALL | Diagnosis |  | Other | JAK2 |
| B-P0195-R1-x | BCP-ALL | Relapse |  | TCF3::HLF | TCF3::HLF |
| B-P0029-D-x | BCP-ALL | Diagnosis |  | TCF3::HLF | TCF3::HLF |
| B-P0030-D-x | BCP-ALL | Diagnosis |  | TCF3::HLF | TCF3::HLF |
| B-P0024-D-x | BCP-ALL | Diagnosis |  | TCF3::HLF | TCF3::HLF |
| B-P0025-D-x | BCP-ALL | Diagnosis |  | TCF3::HLF | TCF3::HLF |
| B-P0485-D-x | BCP-ALL | Diagnosis | Pre-B | TCF3::PBX1 | TCF3::PBX1, TP53 |
| B-P0492-R1-x | BCP-ALL | Relapse | Pre-B | TCF3::PBX1 | TCF3::PBX1 |
| B-P0478-D-x | BCP-ALL | Diagnosis |  | TCF3::PBX1 | TCF3::PBX1 |
| B-P0478-R1-x | BCP-ALL | Relapse |  | TCF3::PBX1 | TCF3::PBX1 |
| B-P0492-R1-x | BCP-ALL | Relapse | Pre-B | TCF3::PBX1 | TCF3::PBX1 |
| B-P1050-R1-x | BCP-ALL | Relapse |  | BCR::ABL1 | BCR::ABL1 |
| B-P0887-R1-x | BCP-ALL | Relapse | Common B | BCR::ABL1 | BCR::ABL1 |
| B-P0887-D-x | BCP-ALL | Diagnosis | Common B | BCR::ABL1 | BCR::ABL1 |
| B-P1329-R1-x | BCP-ALL | Relapse | Common B | BCR::ABL1 | BCR::ABL1, IKZF1 (del) |
| B-P1003D-x | BCP-ALL | Diagnosis |  | KMT2Ar | KMT2A::AFF1 |
| B-P0974-D-x | BCP-ALL | Diagnosis | Pre-B | KMT2Ar | KMT2A::MLLT10 |
| T-P0142D-x | T-ALL | Diagnosis | Pre- T |  |  |
| T-P0356-R1-x | T-ALL | Relapse |  |  |  |
| T-P0360-R1-x | T-ALL | Relapse |  |  |  |
| T-P0967-R1-x | T-ALL | Relapse | Cortical T | Other | CDKN2A (del), CDKN2B (del), NOTCH1, FBXW7 |
| T-P0856-D-x | T-ALL | Diagnosis | Cortical T |  |  |
| T-P1218-D-x | T-ALL | Diagnosis | Pro-T | Other | WT1, NF1, ASXL1, SETD2, BLM, EP300, AFF1, PICALM::MLLT10 |
| T-P0053D-x | T-ALL | Diagnosis | Pre-T |  |  |
| T-P0210-D-x | T-ALL | Diagnosis | Cortical T |  |  |
| T-P0166-D-x | T-ALL | Diagnosis | Cortical T |  |  |
| T-P0070-D-x | T-ALL | Diagnosis | Cortical T |  |  |
